# Self-Assembled Protein Vesicles as Vaccine Delivery Platform to Enhance Antigen-Specific Immune Responses

**DOI:** 10.1101/2024.02.26.582169

**Authors:** Yirui Li, Julie A. Champion

## Abstract

Self-assembling protein nanoparticles are beneficial platforms for enhancing the often weak and short-lived immune responses elicited by subunit vaccines. Their benefits include multivalency, similar sizes as pathogens and control of antigen orientation. Previously, the design, preparation, and characterization of self-assembling protein vesicles presenting fluorescent proteins and enzymes on the particle surface have been reported. Here, a full-size model antigen protein, ovalbumin (OVA), was genetically fused to the recombinant vesicle building blocks and incorporated into protein vesicles via self-assembly. Characterization of OVA protein vesicles showed room temperature stability and tunable size and antigen loading ratio. Immunization of mice with OVA protein vesicles induced strong antigen-specific humoral and cellular immune responses. This work demonstrates the potential of protein vesicles as a modular platform for delivering full-size antigen proteins that can be extended to pathogen antigens to induce antigen specific immune responses.

## 1. Introduction

Vaccines play a vital role as public health interventions against infectious diseases [1, 2]. Traditional vaccines consist of live attenuated or inactivated virus [3, 4]. Although these whole pathogen vaccines elicit strong and long-lasting protective immune response, they do not offer efficient protection against some diseases and are not safe for immunocompromised persons. Protein subunit vaccines, which only include selected antigens from pathogens, offer a safe alternative to live attenuated and inactivated viruses [5, 6]. Moreover, subunit vaccines enable control over antigen-specific immune response, and they are easier to manufacture compared to traditional vaccines. However, subunit vaccines are generally limited by weak and short-lived humoral and cellular immune responses. Therefore, subunit vaccines are often required to be co-administered with adjuvants or in delivery systems for sufficient immune response [7, 8].

In recent decades, nanoparticle delivery systems have been widely studied for subunit vaccine delivery [8-11]. Nanoparticles offer several beneficial features, which include multivalent antigen display to B cell receptors, enhanced uptake by antigen presenting cells, protection of the antigen against degradation, and co-delivery of antigens and adjuvants. Liposomes, polymeric nanoparticles, and protein nanoparticles are commonly investigated for subunit vaccine delivery [12-14]. In liposome and polymeric nanoparticle systems, antigens are either encapsulated within the core or conjugated on the surface. However, both systems face the challenges of low antigen encapsulation and denaturation of antigens during fabrication or chemical conjugation [11, 15, 16]. Virus like particles (VLPs) are the only FDA approved protein nanoparticle for vaccination[17, 18]. As they are made from viral coat proteins, they are quite immunogenic but also introduce off target antigens from the VLP scaffold itself, thereby reducing the control over antigen specificity that is often desired in a subunit vaccine. A number engineered protein nanoparticles have been developed to improve control over both antigen presentation and physical properties[14]. Protein nanoparticles can be formed via desolvation or self-assembly. In the desolvation process, organic solvents may denature antigen proteins, thereby hampering their ability to induce humoral immune responses [19, 20]. In self-assembling protein nanoparticle systems, antigens are genetically fused to protein or peptide building blocks to display antigens on the surface [14, 21]. Self-assembling systems are beneficial for vaccine delivery because they mimic the multivalent, oriented antigen display and the size of natural pathogens (∼20 nm to 400 nm) while preserving the selectivity of subunit vaccines. Furthermore, self-assembly in aqueous buffer better maintains the structure of antigen proteins. Self-assembling peptide cages (SAGEs) and self-assembling protein nanoparticle (SAPNs) are two examples of self-assembling systems for subunit vaccine delivery [22-26]. Both systems have shown efficacy to provide protection against infectious disease in animal models. However, only antigenic peptides and very small proteins, such as 32 amino acid Helix C of influenza, were genetically fused to their building blocks instead of full-sized, folded antigen proteins, as genetic fusion to larger antigen proteins may hamper self-assembly. Large protein antigens can provide broad-spectrum protection by eliciting immune responses against multiple epitopes with conformations that are identical to conformations in pathogens [27, 28]. *De novo* designed icosahedral protein cages that resemble VLPs, but have no viral components, are capable of displaying large protein antigens such as DS-Cav1 from Respiratory syncytial virus and SARS-CoV-2 spike [29, 30]. Additionally, a variety of natural protein cages such as human ferritin, can also be recombinantly fused to antigens, including SARS-CoV-2 spike [14, 31].

In addition to SAGEs, SAPNs and protein cages, self-assembling elastin-like polypeptides (ELPs) have been explored for designing nanoparticle vaccines. ELP is a thermosensitive peptide with the typical sequence (VPGXG)_n_ that undergoes a hydrophobic transition upon warming that leads to coacervation and phase separation[32-34]. ELP-based nanoparticle vaccines were self-assembled from ELPs that were genetically fused to antigenic peptides [35, 36] or small antigen protein, M2e (23 amino acids) [37]. No ELP-based nanoparticle that delivers full-size antigen proteins has been reported yet. Our group has previously developed protein vesicles self-assembled from full-size, globular proteins, including fluorescent proteins [38] and enzymes [39]. The proteins are fused to acidic leucine zipper, Z_E_, and mixed with ELP fused to basic leucine zipper, Z_R_ (Z_R_-ELP). Z_E_ and Z_R_ bind each other with high (10^15^ M) affinity [40]. Upon warming, the proteins transition from soluble to unstable coacervates to stable, hollow vesicles [41]. Furthermore, protein vesicle diameter and membrane structure could be tuned by varying molar ratio of Z_E_ to Z_R_ [38], protein concentration [41], and NaCl concentration [42]. Inspired by these protein vesicles that display large globular proteins on their surface, we explored the potential of protein vesicles as a novel vaccine platform to deliver antigen proteins. In this study, we chose a model antigen protein, ovalbumin (OVA), which is known to be immunogenic and has been commonly used to investigate the efficacy of nanoparticle vaccine platform [43-45]. OVA was genetically fused to Z_E_ to incorporate OVA into protein vesicles through self-assembly. Non-natural amino acid, para-azidophenylalanine (pAzF) was incorporated into Z_R_-ELP (pZ_R_-ELP) to enable photo-crosslinking of OVA protein vesicles. To examine the efficacy of protein vesicles for vaccine delivery, we characterized the self-assembly and nanostructure of OVA protein vesicles and performed *in vitro* experiments and *in vivo* vaccination.

## 2. Materials and Methods

### 2.1 Expression and Purification of Proteins

pET17b-OVA-Z_E_ plasmid was purchased from Genscript. *E. coli* Shuffle T7 strain (NEB) was transformed with pET17b OVA-Z_E_. To express OVA-Z_E_, a single colony was inoculated in 10 mL lysogeny broth (LB) medium containing 200 mg/L ampicillin and shaken overnight at 30 °C. 30 mL overnight culture was inoculated in 1 L LB media. When optical density at 600 nm (OD600) reached ∼0.8 to 1, 1 mM isopropyl-β-thiogalactoside (IPTG) was added to induce protein expression, and the cultures were shaken at 16 °C for an additional 18 hours. Cells were harvested by centrifugation (4000 g, 10 minutes) and stored at -20 °C. The pellets were resuspended in lysis buffer (300 mM NaCl, 50 mM NaH_2_PO4, and 10 mM imidazole), followed by sonication for 15 mins. After centrifugation (10000 g, 30 minutes), the supernatant was incubated with Ni-nitrilotriacetic acid (NTA) agarose resin (Qiagen) for 1 hour at 4 °C. The suspension was flowed through an Econo-Column (Biorad) and washed with 100 mL of lysis buffer containing 40 mM imidazole. Elutions were collected using buffer with 250 mM imidazole. The protein elutions were buffer exchanged into phosphate buffered saline (PBS) by dialysis with four buffer exchanges at 4 °C.

pZ_R_-ELP was produced as described previously [42]. *E. coli* strain AFIQ-BL21[46] was transformed with pQE60 Z_R_-ELP containing a mutant *E. coli* phenyl-alanyl-tRNA synthetase (A294G) gene [47, 48]. Briefly, a single colony was grown overnight and inoculated in 1 L M9 minimum media supplemented with glucose (0.4 wt %), thiamine (5 mg/L), MgSO_4_ (1 mM), CaCl_2_ (0.1 mM), and 20 natural amino acids at a concentration of 1 mg/L each. At OD600 of 0.8, cells were harvested by centrifugation and resuspended in M9 minimum media supplemented with 19 natural amino acids at 1 mg/L each except phenylalanine and 0.3 mg/L pAzF (Bachem). After 15 minutes, 1 mM IPTG was added for induction. After 5 h, cells were harvested by centrifugation and purified with Ni-NTA resin using non-native buffers as described previously [38]. Purified pZ_R_-ELP was dialyzed (3k MWCO) into Milli-Q water and lyophilized. All protein sample containers were wrapped in foil and kept in the dark to protect pAzF from ambient light. Protein purity was verified by sodium dodecyl sulfate polyacrylamide gel electrophoresis (SDS-PAGE).

### 2.2 Vesicle Assembly and Turbidity Measurement

Protein solutions were prepared on ice by adding water, pZ_R_-ELP, and OVA-Z_E_, then adding 10X PBS to achieve a specified salt concentration based on NaCl in 10X PBS. Solutions were moved to the bench (25 °C) for 1 hour to induce vesicle assembly. Vesicles were crosslinked by UV irradiation at 254 nm for 30 minutes. The turbidity of protein solutions was measured at OD 400 nm, using a microplate reader (Synergy HT Multi-Mode, BioTek). 100 μL of protein solutions were prepared in a 96-well microplate at 4 °C and placed in the microplate reader at 25 °C. Then, the changes of turbidity were monitored by recording the OD of protein solutions every minute for 1 hour.

### 2.3 Dynamic Light Scattering (DLS)

Hydrodynamic diameters of protein vesicles were measured by Zetasizer DLS instrument (Malvern Instruments, Malvern). A 4 mW He-Ne laser operating at a wavelength of 633 nm was equipped and operated at a detection angle of 173°. 70 μL of vesicle solution was prepared in a microcuvette and measured at 25 °C. The solvent was adjusted to aqueous solutions containing 0.50 M to 1.5 M NaCl buffered with phosphate and the material was protein. Z-average values were used to report vesicle size.

### 2.4 Transmission Electron Microscopy (TEM)

OVA protein vesicles were imaged by a TEM (JEM 100CX-II, JEOL). 5 μL of protein vesicle solution was dropped on a copper grid (Electron Microscopy Sciences) for 5 min at 25 °C, washed with Milli-Q water, stained with 1% phosphotungstic acid solution for 20 s, and washed with Milli-Q water. TEM samples were air-dried for 24 hour and imaged at 100 kV.

### 2.5 Circular Dichroism (CD)

The CD spectra of soluble OVA-Z_E_ and photo-crosslinked OVA protein vesicles were obtained by Chirascan-plus CD spectrometer (Applied Photophysics). Measurements were performed in a 0.2 cm length cuvette at 25 °C. The spectra were obtained in 1 nm increments within a wavelength range of 200−280 nm.

### 2.6 Endotoxin Removal

Endotoxin levels in OVA-Z_E_ and pZ_R_-ELP were reduced by Pierce™ High Capacity Endotoxin Removal Resin (Thermo Fisher). To confirm low endotoxin content within proteins, the endotoxin levels in OVA-Z_E_ and pZ_R_-ELP were quantified by ToxinSensor™ Chromogenic LAL Endotoxin Assay Kit (GenScript). OVA-Z_E_ and pZ_R_-ELP with low endotoxin levels and endotoxin free water and PBS were used to prepare OVA protein vesicles for *in vitro* and *in vivo* studies. The endotoxin level in all samples was less than 5 EU/kg as recommended by the United States Pharmacopoeia [49].

### 2.7 *In Vitro* Dendritic Cell Maturation

The JAWS II immature dendritic cell (DC) line (ATCC) was cultured in MEM-alpha (Corning) supplemented to 4 mM glutamine and 5 ng/mL GM-CSF (Peprotech), 20% fetal bovine serum (FBS), and 1% penicillin/ streptomycin (Amresco). JAWS II DCs were plated in 48-well plates for measuring *in vitro* maturation. Cells were stimulated with 5 μg/mL of bacterial lipopolysaccharide (LPS) and 20 μg/mL of soluble OVA, soluble OVA-Z_E_ or OVA protein vesicles. After 24 hours, cells were fixed with 3.7% paraformaldehyde and blocked with 1% BSA in PBS for 1 hour. Next, cells were incubated with 1 μg/μL of TruStain FcX antibody (BioLegend) for 10 minutes on ice to block non-specific binding to Fc receptors. After that, cells were stained with 1 μg/μL of PE anti-mouse CD86 (BioLegend) for 30 minutes. Finally, cells were washed twice with PBS and scraped from the plate. The upregulation of CD 86 was analyzed by flow cytometry.

### 2.8 Immunization of mice

All animals were treated in accordance with the regulations and guidelines of the NIH Guide for the Care and Use of Laboratory Animals, and all protocols and procedures were reviewed and approved by Georgia Tech’s Institutional Animal Care and Use Committee (A100529). 6-8 week-old Balb/c mice (Jackson Laboratory) were immunized intramuscularly at the right back leg with solutions containing 10-12 μg of soluble or vesicle form of OVA-Z_E_ in pharmaceutical grade saline. Prime vaccination was given at day 0 and booster vaccination was given at day 21. Animals were monitored for weight loss and signs of lethargy after vaccination for 7 days.

### 2.9 Blood collection

Approximately 100 μL of blood was collected from immunized mice by jugular vein puncture immediately prior to vaccination and 2 weeks after prime and boost vaccinations. Blood was allowed to clot in BD Vacutainer tubes (Becton, Dickinson & Company) and was centrifuged at 1000 g for 15 minutes to collect serum. Serum was stored at -20 °C.

### 2.10 Antibody Endpoint Titer Measurement

Nunc MaxiSorp plates (Thermo Fisher) were coated with 1 μg/mL of purchased OVA (Sigma) in PBS overnight at room temperature. Plates were blocked with 1% BSA in PBS-Tween 20 (PBST) for 1 hour followed by three washes with PBST. Next, serum was added at a 1/10 dilution and subsequent 5-fold serial dilutions in 1% BSA/PBST and incubated for 1 hour at room temperature. Serum from naïve mice (pre-vaccination) was also run on each plate to determine cutoff values. After serum incubation, plates were washed with PBST and incubated with horseradish peroxidase (HRP)-conjugated goat anti-mouse IgG1 antibodies (Southern Biotech) or HRP-conjugated goat anti-mouse IgG2a-HRP antibodies (Southern Biotech) at a 1:4000 dilution in 0.1% BSA/PBST for 1 hour at room temperature followed by washing with PBST. 100 μL TMB chromogen solution (Thermo Fisher) was added into each well. After 30 minutes, the enzymatic reaction was quenched with H_2_SO_4_ (Thermo Fisher). OD was read at 450 nm using a microplate reader (BioTek). End point titers were determined from reciprocal dilutions to determine the dilution at which the OD value was equal to the mean + three standard deviations of that of naïve serum [50]. Titer values too low for detection were fixed at 10, corresponding to the lowest dilution used.

### 2.11 Preparation of Splenocytes and Lymphocytes

2 weeks after boost immunization, mice were sacrificed by euthanasia, and spleens were collected. Spleens were manually homogenized by the plunger of a 3 ml syringe (Becton, Dickinson & Company) in a 100 μM cell strainer (Greiner Bio-One). Single cell suspensions of splenocytes were prepared by forcing cells through cell strainers with complete RPMI media (RPMI, 2mM L-Glutamine, 50 μM 2-mercaptoethanol, 100 U/mL penicillin, 100 μg/mL streptomycin, and 10 % FBS). Cells were centrifuged at 350 g for 5 minutes. Splenocytes were resuspend in 1 mL ACK lysing buffer (Thermo Fisher) to lyse red blood cells. After 10 minutes, splenocytes were centrifuged and resuspended in complete RPMI media.

### 2.12 Intracellular Cytokine Staining

Splenocytes were plated in 96-well U-bottom plates at 10^6^ cells/well in RPMI and stimulated with 1 μg/mL of PepTivator Ovalbumin (Miltenyi Biotec). After 3 hours, each well was supplemented with a transport inhibitor, 1X brefeldin A (Biolegend), and incubated for an additional 3 hours. Cells in each well were centrifuged at 350 g for 5 minutes and resuspended in 100 μL of PBS premixed with 0.5 μL of Trustain FcX plus blocking solution (BioLegend) on ice for 10 min. Cells were centrifuged and stained by 100 μL of antibody cocktail solution containing Zombie Violet, PerCP anti-mouse CD3ε antibody, FITC anti-mouse CD8a antibody and APC/Cyanine7 anti-mouse CD4 antibody (BioLegend) in PBS. After staining for 1 hour in the dark, cells were washed with 1% BSA in PBS, centrifuged, and fixed with 100 μL of 3.7% formaldehyde (VWR) in PBS for 30 min on ice. After fixation, cells were centrifuged and washed once with 100 μL of permeabilization buffer (eBioscience). Cells were centrifuged and resuspended in 100 μL of permeabilization buffer containing 1.5 μL of PE anti-mouse IFN-γ antibody and 1.5 μL of PE/Cyanine7 anti-mouse IL-4 antibody. After incubation on ice for 30 minutes, cells were centrifuged, washed with 1% BSA in PBS, and resuspended in 100 μL of 1% BSA in PBS. Flow cytometry analysis was conducted using Cytek Aurora (Cytek Biosciences). Data was analyzed with Flow Jo (Becton, Dickinson & Company).

### 2.13 Statistical Analysis

Antibody endpoint titers were analyzed using the Mann–Whitney U test and T cell counts were analyzed by one-way analysis of variance (ANOVA) with Tukey’s post-hoc multiple comparison. All statistical analysis were conducted using GraphPad Prism 9 (GraphPad). The p values < 0.05 were considered statistically significant (*p < 0.05, **p < 0.01).

## 3. Results and Discussion

### 3.1 Synthesis of recombinant OVA-Z_E_ and pZ_R_-ELP

OVA-Z_E_ was expressed in the Shuffle T7 express strain, which is an engineered *E. coli* strain to promote disulfide bond formation in the cytoplasm [51], and the final yield was ∼1 mg/L of *E*.*coli* culture (Figure S1a and S1b). To verify that fully folded OVA was produced, we compared the secondary structure of recombinant OVA-Z_E_ with native OVA measured by circular dichroism (CD). According to the CD spectra (Figure S1c), OVA-Z_E_ showed similar, but slightly different structure from native OVA, likely due to the fusion to Z_E_ motif and lack of post-translational modification. As *E. coli* expression does not provide native post-translational modifications to viral (or egg) proteins, and it is challenging to express antigens with complex structures. Furthermore, endotoxin contamination requires extra processing steps after antigen production. Therefore, insect cells or mammalian cells can be used for antigen protein expression in future work, as is common in subunit viral vaccines [52]. The second component of vesicles is Z_R_-ELP. To stabilize protein vesicles, a photo-crosslinkable non-natural amino acid, pAzF, was incorporated into phenylalanine residues in the ELP domain by the global incorporation method using phenylalanine auxotroph AFIQ *E. coli* containing a mutant phenylalanyl-tRNA synthetase, as described previously [42, 48, 53]. UV crosslinking creates covalent bonds between the azide group and nearby protein backbone, as evidenced by oligomers of pZ_R_-ELP seen in SDS-PAGE (Figure S2).

### 3.2 Tunable size and antigen loading of OVA protein vesicles

To prepare OVA protein vesicles, 0.3 – 4.5 μM OVA-Z_E_ and 30 μM pZ_R_-ELP were mixed in PBS buffer containing 0.5 – 1.0 M NaCl at 4 °C. Increasing the temperature to 25 °C triggered the ELP phase transition and OVA-Z_E_/pZ_R_-ELP complexes self-assembled into OVA protein vesicles (Figure 1). The turbidity profiles of protein mixtures during phase transition were measured (Figure S3). The initial rapid increase indicated occurrence of the ELP phase transition in the protein mixtures and self-assembly and growth of particulate like coacervates. Once the turbidity profile reached saturation it stabilized, indicating that the coacervates reorganized so that the hydrophilic OVA-Z_E_ shielded the hydrophobic ELP interior, resulting in the formation of stable OVA protein vesicles.

**Figure 1.**
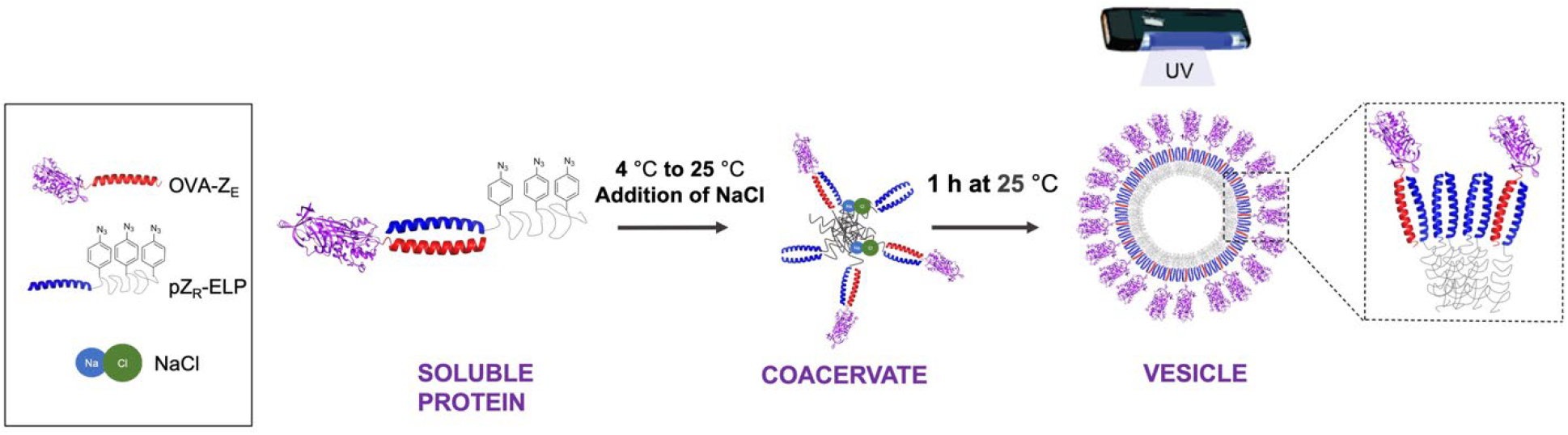
Schematic of thermally triggered self-assembly of OVA protein vesicles. OVA-Z_E_ and pZ_R_-ELP were mixed in phosphate buffer containing additional NaCl at 4 °C. Protein mixtures were then incubated at 25 °C for 1 hour for vesicle self-assembly and crosslinked by UV irradiation at 254 nm for 30 minutes.

Previous work on protein vesicle has shown that the sizes of protein vesicles could be tuned by changing Z_E_/Z_R_ molar ratios [38] or NaCl concentrations [42]. To optimize OVA protein vesicles for vaccination, DLS was used to measure hydrodynamic diameter of vesicles formed at different Z_E_/Z_R_ molar ratios and NaCl concentrations. Increasing the Z_E_/Z_R_ molar ratio, at constant pZ_R_-ELP and NaCl concentration, led to changes in the molecular packing in protein amphiphiles. A greater number of hydrophilic OVA proteins increased the curvature of protein vesicles to accommodate steric hinderance between OVA (Figure 2a). This effect saturates and we have observed previously that increasing the molar ratio beyond a critical level results in a population of soluble globular proteins co-existing with vesicles [54], though no evidence for soluble protein was observed in this work. Increasing NaCl concentration, at constant protein concentration, increases ELP hydrophobicity [55], resulting in more compact ELP conformation. This, too, leads to decreased vesicle sizes as the relative size of OVA to ELP increases and increased curvature is required to relieve steric constraints of OVA packed closer together due to smaller interior ELP domains (Figure 2b). Altogether, this data is consistent with previous data on protein vesicles made with other globular proteins and confirms that vesicle size can be tuned. The ability to tune the density of antigen proteins on the vesicle surface is highly relevant to vaccine design and future work could explore how antigen density can be best matched to B cell receptor binding for the greatest antibody response. As lower antigen densities on vaccine nanoparticles have been reported to be better than high densities [56], we selected nanoscale vesicles made at 0.1 Z_E_/Z_R_ molar ratio and 1.0 M NaCl for further characterization and vaccination.

**Figure 2.**
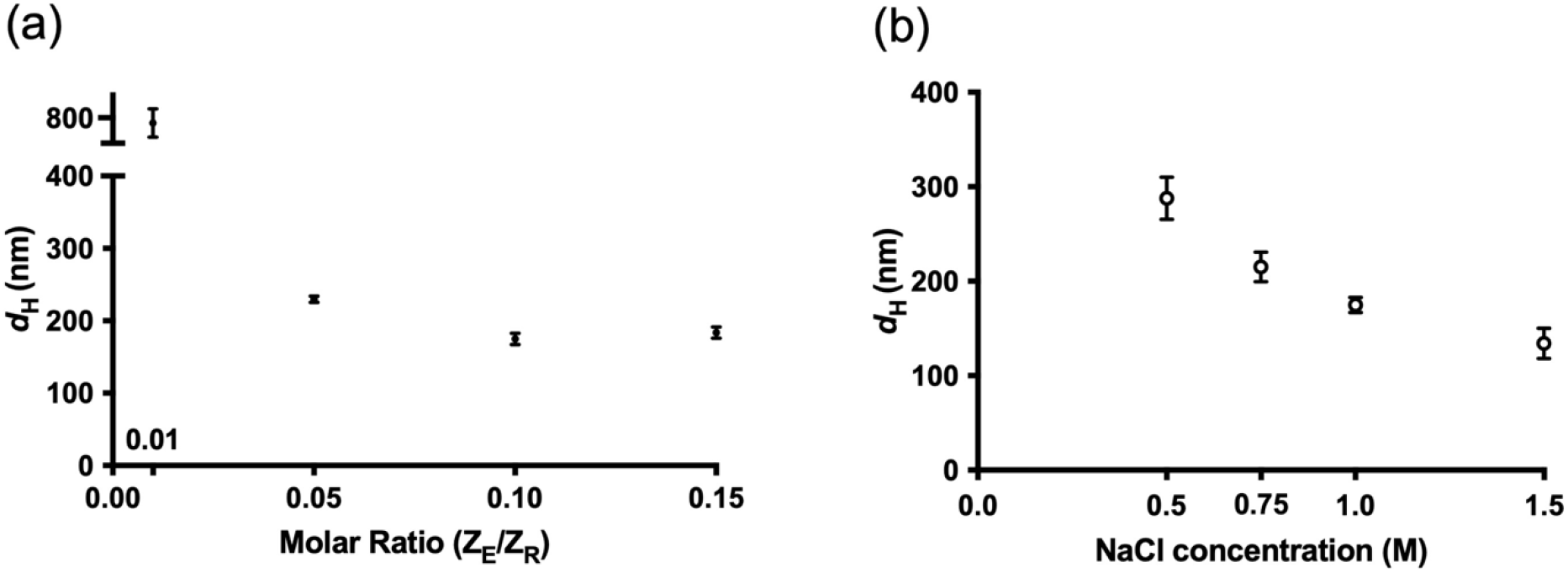
Synthesis of OVA protein vesicles at various molar ratio and NaCl concentrations. (a) Hydrodynamic diameters of OVA protein vesicles formed at various Z_E_/Z_R_ molar ratios from 30 μM pZ_R_-ELP and 1.0 M NaCl. (b) Hydrodynamic diameters of OVA protein vesicles formed at various NaCl concentrations with 30 μM pZ_R_-ELP and 0.1 Z_E_/Z_R_ molar ratio. Each data point is the average of three replicate batches of vesicles.

### 3.3 OVA protein vesicle characterization

Vesicles made from 30 μM pZ_R_-ELP, 3 μM OVA-Z_E_, and 1.0 M NaCl were crosslinked with UV irradiation at 254 nm for 30 minutes. This is necessary because the ELP phase transition is reversible. We previously showed that dilution of similar vesicles made with mCherry-Z_E_ and pZ_R_-ELP at 1.0 M salt into physiological salt (0.15 M) resulted in swelling and disassembly of the vesicles [42]. The vesicles exhibited a typical turbidity profile (Figure S4) and a hydrodynamic diameter of 174.9 ± 8 nm with a polydispersity index of 0.2 ± 0.03 (Figure 3a). TEM of OVA protein vesicles showed spherical shapes with wrinkled surface, indicative of hollow vesicle structures (Figure 3b). To avoid hypertonic shock during *in vitro* and *in vivo* experiments, OVA protein vesicles initially formed in 1 M NaCl were dialyzed into PBS to match physiological salt concentration. Diameter of vesicles were monitored for 7 days after dialysis. OVA protein vesicles showed initial swelling to 271 nm at day 2 and were stable in PBS for 7 days (Figure S5). Furthermore, we kept vesicles at room temperature for 44 days. Size distributions of the same sample on day 1 and day 44 proved that protein vesicles maintain their structures at room temperature at least 44 days (Figure 3a). The long-term stability of protein vesicles at room temperature is beneficial as it can eliminate the need for high-cost cold chain shipping and storage that makes vaccines inaccessible for resource-limited regions [57, 58]. To confirm UV irradiation did not denature antigen proteins on the surface, we compared the secondary structure of OVA protein vesicles with and without UV irradiation measured by CD. OVA protein vesicles with and without UV irradiation displayed comparable CD spectra (Figure S6), which proved that UV irradiation did not damage the structure OVA antigens on the surface. Some antigens, however, may be susceptible to denaturation by UV irradiation. In this case, the amino acid sequence of the ELP domain in Z_R_-ELP can be tuned to increase the hydrophobicity and enable nanoscale vesicle assembly and stability at physiological salt concentration, which we have recently demonstrated [59].

**Figure 3.**
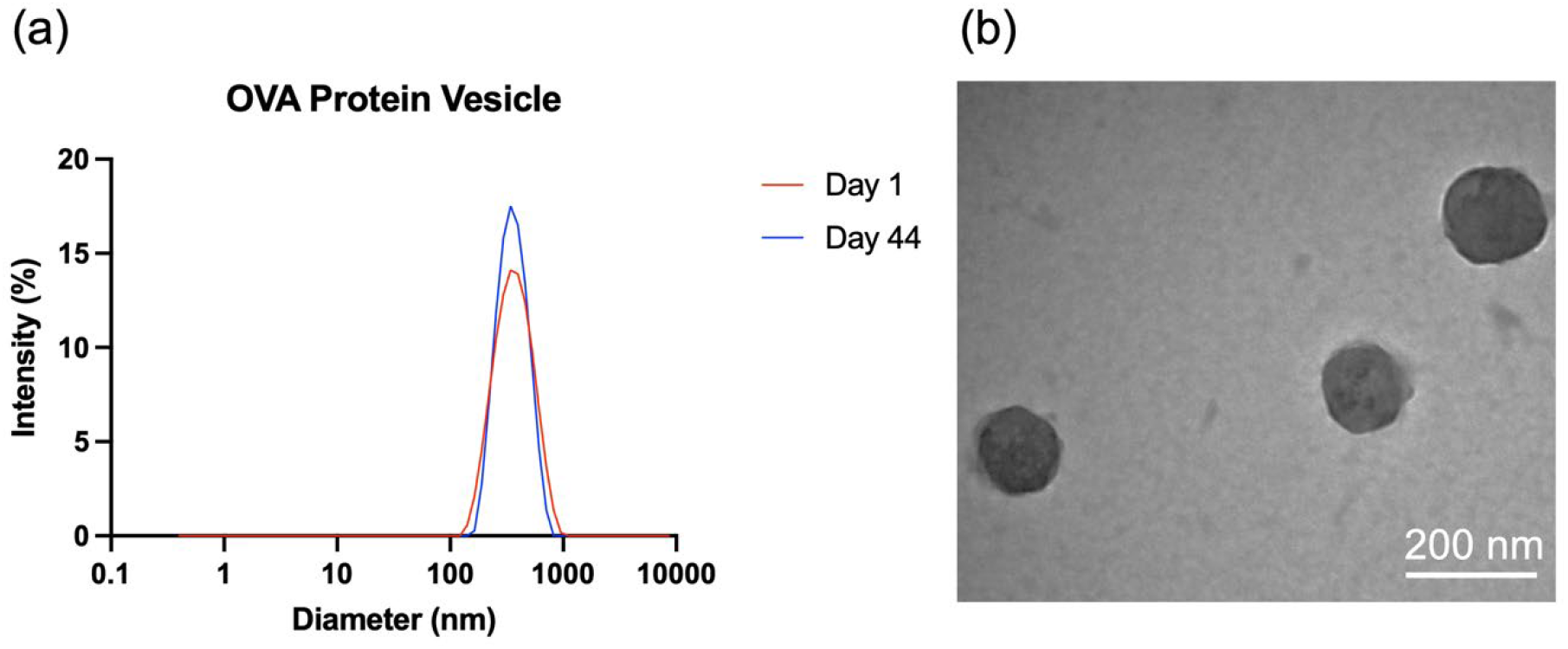
Self-assembly of recombinant OVA-Z_E_ and pZ_R_-ELP into OVA protein vesicles. (a) Size distributions of OVA protein vesicles in PBS at day 1 (red) and day 44 (blue). (b) TEM micrograph of OVA protein vesicles. Scale bar = 200 nm.

Similar to SAGEs, SAPNs, and protein cages, OVA protein vesicles mimic the repetitive antigen display and particulate feature of natural pathogens [22-25]. However, SAGEs and SAPNs only showed the ability to present antigenic peptides or very small (∼30 amino acids) protein on the surface. It has been challenging to genetically fuse large antigen proteins to engineered self-assembling building blocks as it can cause structural distortion in the final self-assembly [60]. So far, a few self-assembling protein nanoparticle systems have been reported to display full-size antigens [29, 61-63]. The protein vesicle design allows presentation of full-size OVA antigens (385 amino acids) via high affinity Z_E_/Z_R_interactions during self-assembly. Furthermore, our group has demonstrated that two Z_E_ fusion proteins, mCherry-Z_E_and GFP-Z_E_, were incorporated into the same protein vesicle [38]. This feature might be beneficial for designing multiple antigen-presenting vaccines against diseases that require vaccination with multiple antigens for comprehensive immune responses [64, 65]. Matching the natural oligomeric state of some antigens is ideal for designing vaccines with enhanced immune responses, such as trimeric influenza hemagglutinin antigen [66]. Thus, trimeric coiled coils can be explored in future work as building blocks to incorporate trimeric antigens, for example [67].

### 3.4 *In vitro* DC maturation

Dendritic cells (DCs) play an important role as antigen presenting cells (APCs) in the early phase of adaptive immune response [68, 69]. DCs are capable of recognizing pathogens through pattern recognition receptors such as toll-like receptors (TLR). After antigen uptake by DCs, upregulation of maturation markers, including CD86, is necessary for DC maturation and subsequent antigen presentation. To investigate the effect of OVA protein vesicles on JAWS II DC maturation, CD86 was labeled by fluorescent anti-CD86 antibody and upregulation of CD86 was measured by flow cytometry after 24-hour incubation with soluble native OVA, soluble OVA-Z_E_and OVA protein vesicles [43]. Higher CD86 fluorescence in the OVA protein vesicle group indicated that OVA protein vesicles triggered the upregulation of CD86 significantly more than soluble native OVA and OVA-Z_E_(Figure 4). It has been reported that particulate nature and multivalent antigen display facilitates antigen uptake by APCs [70, 71]. Following antigen uptake and DC maturation, antigens are presented via major histocompatibility complex (MHC) molecules to induce T cell activation. Improved antigen presentation by OVA protein vesicles will promote the activation of downstream humoral and cellular immune response and motivates assessment of vesicles by *in vivo* vaccination.

**Figure 4.**
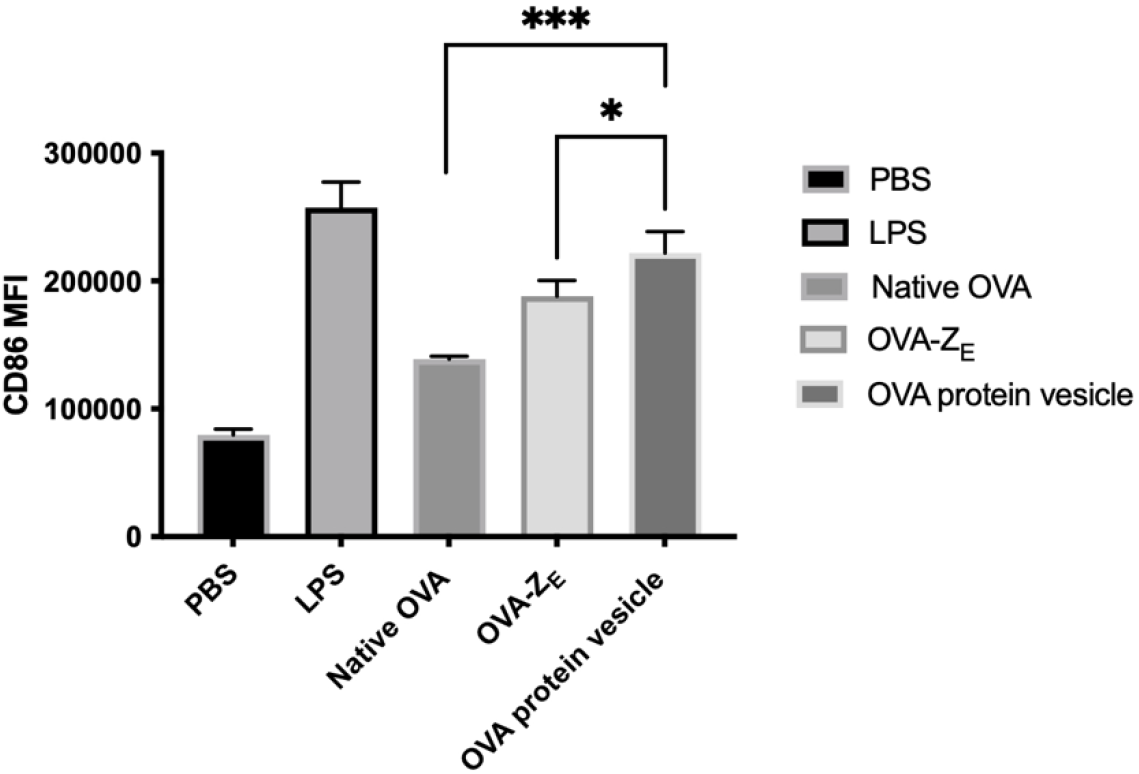
CD86 upregulation in JAWS II DCs after incubation with soluble native OVA, soluble OVA-Z_E_ and OVA protein vesicles for 24 hours. LPS was used as a positive control. Each data point is the average of three replicates of the mean fluorescent intensity (MFI) of the cell population labeled with fluorescent anti-CD86. (***p<0.005, *p<0.05)

### 3.5 OVA-Specific Antibody Responses *In Vivo*

To assess the immune response against OVA protein vesicles, BALB/c mice were immunized intramuscularly with OVA protein vesicles and soluble OVA-Z_E_ in formulations containing 10 μg OVA-Z_E_. The vesicle group also contained 33.8 μg pZ_R_-ELP. A boost immunization of the same formulations was given on day 21 (Figure 5). Blood samples were collected 2 weeks after prime and boost immunizations and OVA-specific antibody endpoint titers were measured by ELISA. After prime immunization, OVA protein vesicles elicited OVA-specific IgG1 and IgG2a antibody responses, while no detectable antibody response was observed in the soluble group (Figure 6). Post boost immunization, IgG1 titers increased for both soluble and vesicle groups. However, OVA protein vesicles still showed ∼20-fold higher IgG1 titer than soluble OVA-Z_E_. IgG2a titers also increased in vesicle vaccinated animals after boost. No detectable IgG2a antibody response was detected in the soluble group even after boost administration. Furthermore, it is known that the multivalent antigen display on nanoparticle surfaces enhances cross-linking between B-cell surface receptors, which favors the production of antibody responses [21, 72]. Therefore, it is likely that protein vesicles presenting multiple OVA antigens on the surface were advantageous for efficient binding and activation of B-cell receptors, resulting in increased antibody titers.

**Figure 5.**
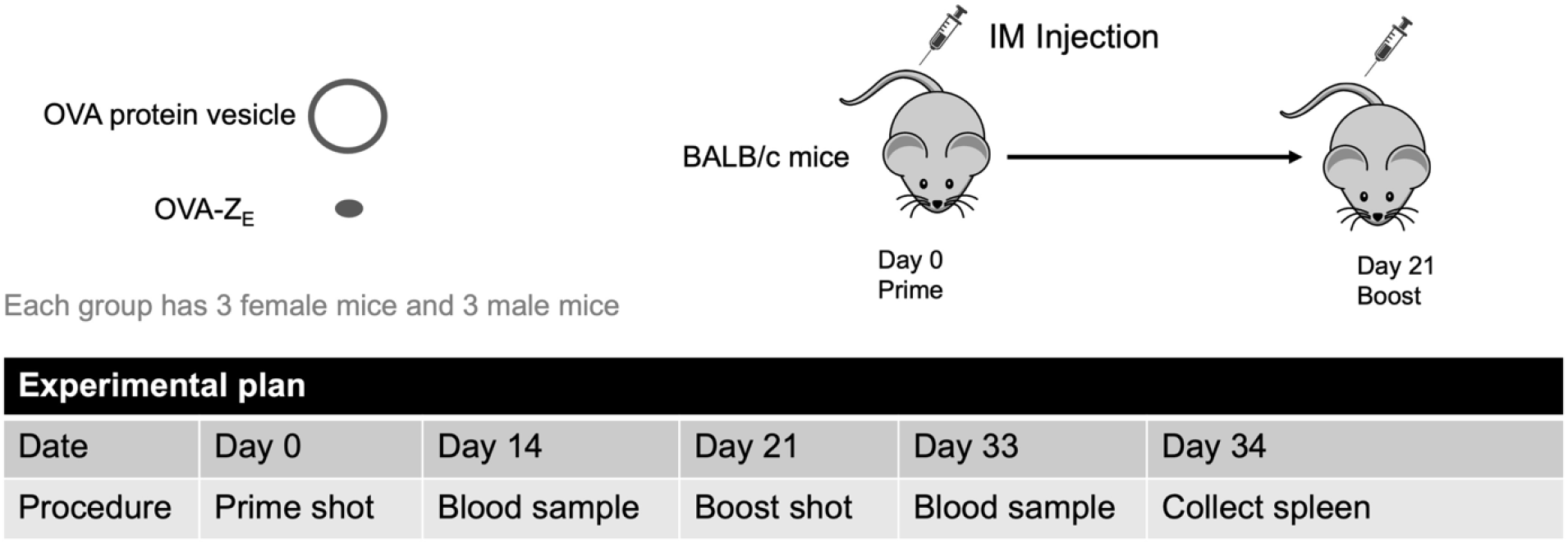
Experimental plan of immunization and sample collection.

**Figure 6.**
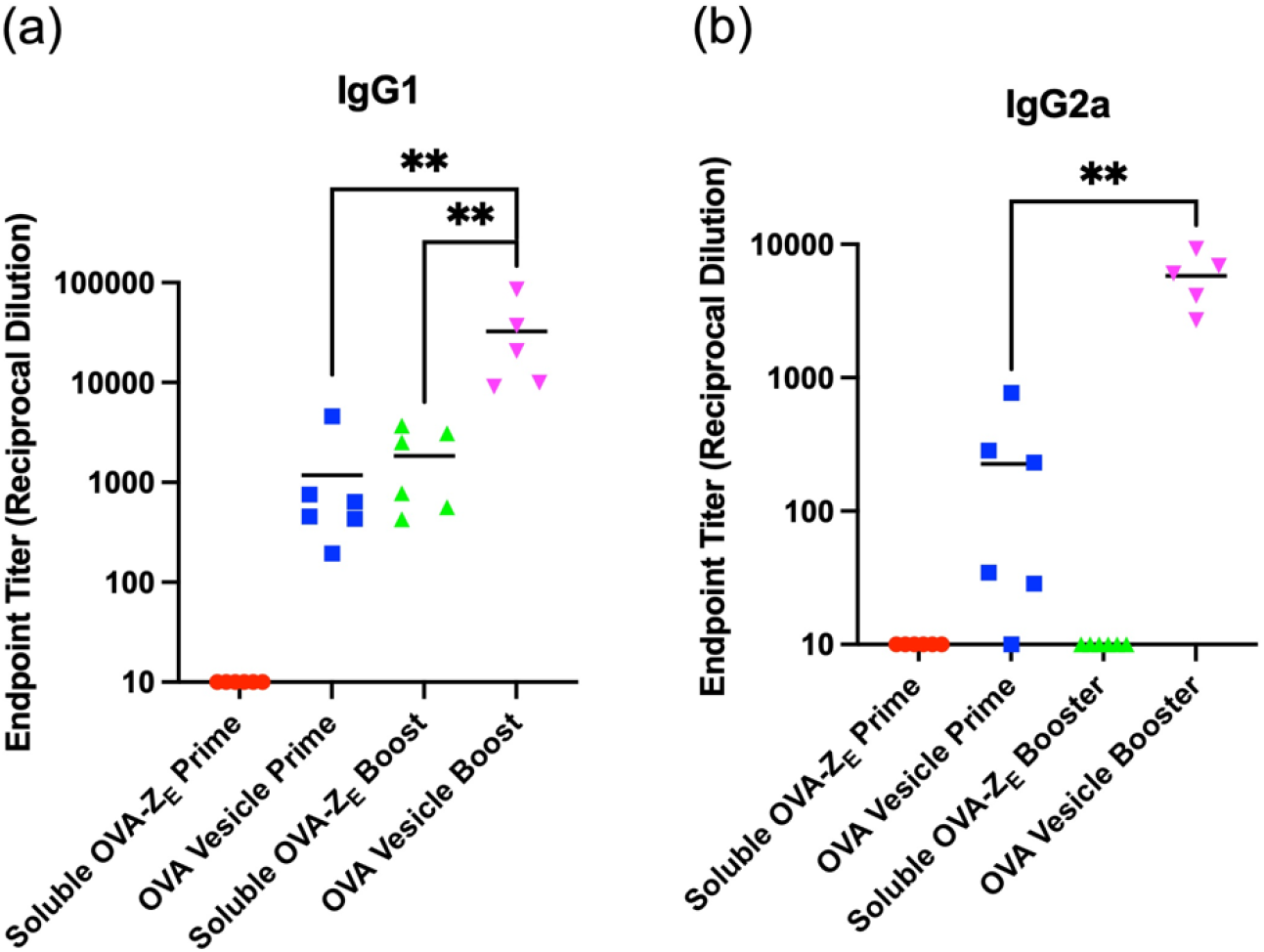
ELISA endpoint titers of OVA specific IgG1 (a) and IgG2a (b) antibodies after prime and boost immunization. Each mouse was immunized with 10 μg OVA-Z_E_ in both soluble OVA-Z_E_ group and OVA protein vesicle group. Titer values too low for detection were arbitrarily set at 10. (**p<0.01)

Though ELPs have been proved to be immunotolerant [73-75], pZ_R_-ELP in nanoparticle form may enhance immune responses against pZ_R_-ELP and neutralizing antibodies against vesicles themselves may hamper the efficiency of vaccine after repetitive vaccination. Therefore, antibody responses against pZ_R_-ELP in OVA protein vesicles were also analyzed in this study. IgG1 antibody against pZ_R_-ELP was detected after prime and boost immunization and IgG2a antibody response was detected after boost immunization (Figure 7). Though pZ_R_-ELP (33.8 μg) was given at a higher dose than OVA-Z_E_ (10 μg), both IgG1 and IgG2a antibody responses against OVA were significantly higher than against pZ_R_-ELP. As OVA antigens were presented on the surface and pZ_R_-ELP proteins were shielded by OVA-Z_E_and embedded inside of the protein vesicles, OVA had higher chance to be processed and presented by APCs than pZ_R_-ELP. Additionally, in a study by Cho et al., immune-tolerant ELPs were designed using atypical ELP sequences derived from homologous mouse and human elastin sequences [76]. As a result, ELPs themselves were not immunogenic in mice while ELP fused with OVA peptide enhanced antibody titers and cytotoxic T lymphocyte responses. If the immunogenicity of pZ_R_-ELP needs to be diminished in future work, the natural sequence from mouse tropoelastin and human elastin can be employed to further reduce antibody responses against pZ_R_-ELP. The fact that vesicle vaccine boost improved anti-OVA titers suggests that neutralization from the low titers of anti- pZ_R_-ELP is not significant. Additionally, a recent report comparing antigen and scaffold antibody responses of *de novo* designed two-component protein cages found that anti-scaffold antibodies do not negatively correlate with anti-antigen responses for a panel of common viral antigens, except subdominant HIV-1 Env antigen [77].

**Figure 7.**
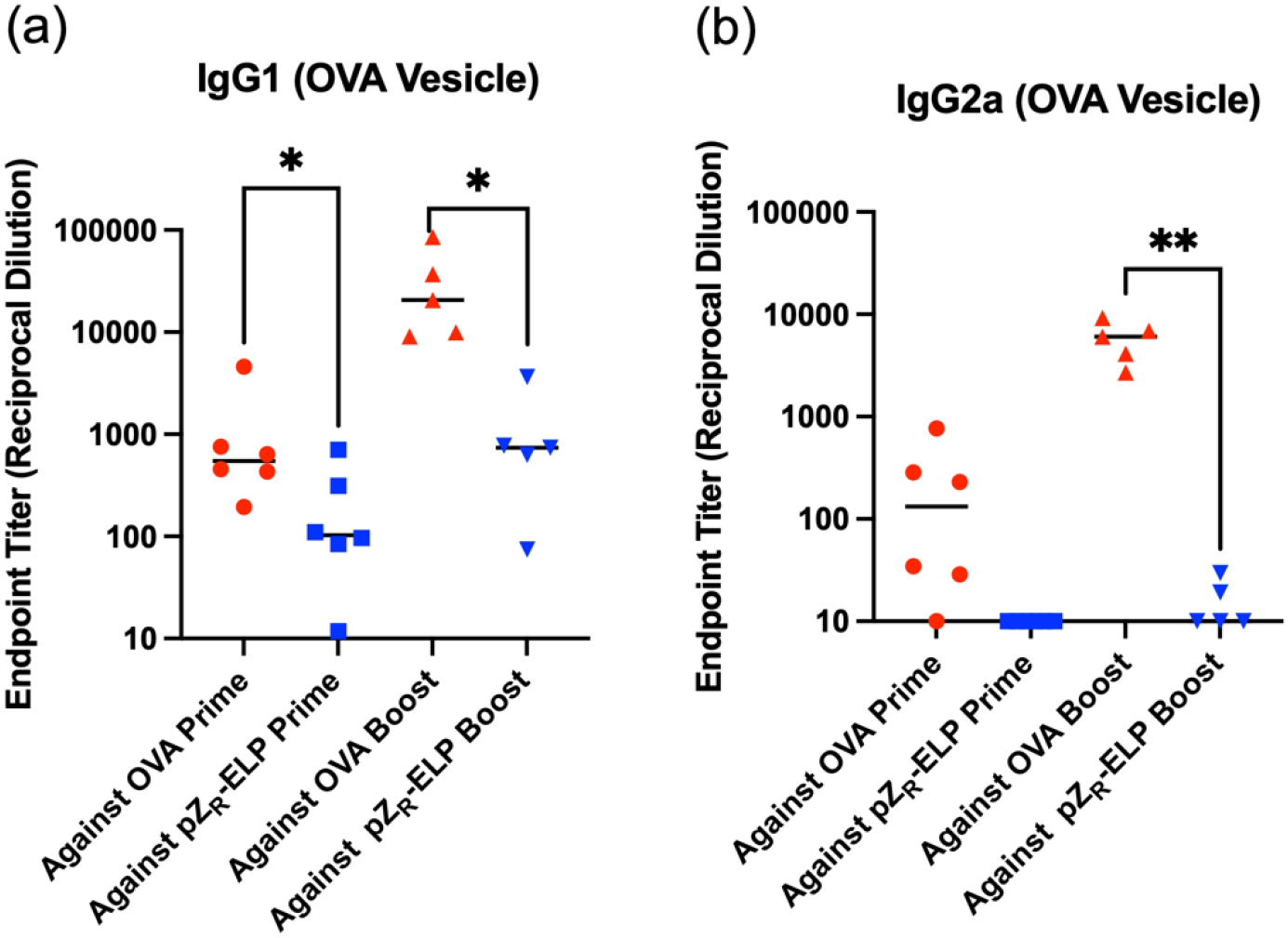
Comparison of ELISA endpoint titers of IgG1 (a) and IgG2a (b) against OVA antigen and pZ_R_-ELP antigen. Titer values too low for detection were arbitrarily set at 10. (**p<0.01, *p<0.05)

**Figure 8.**
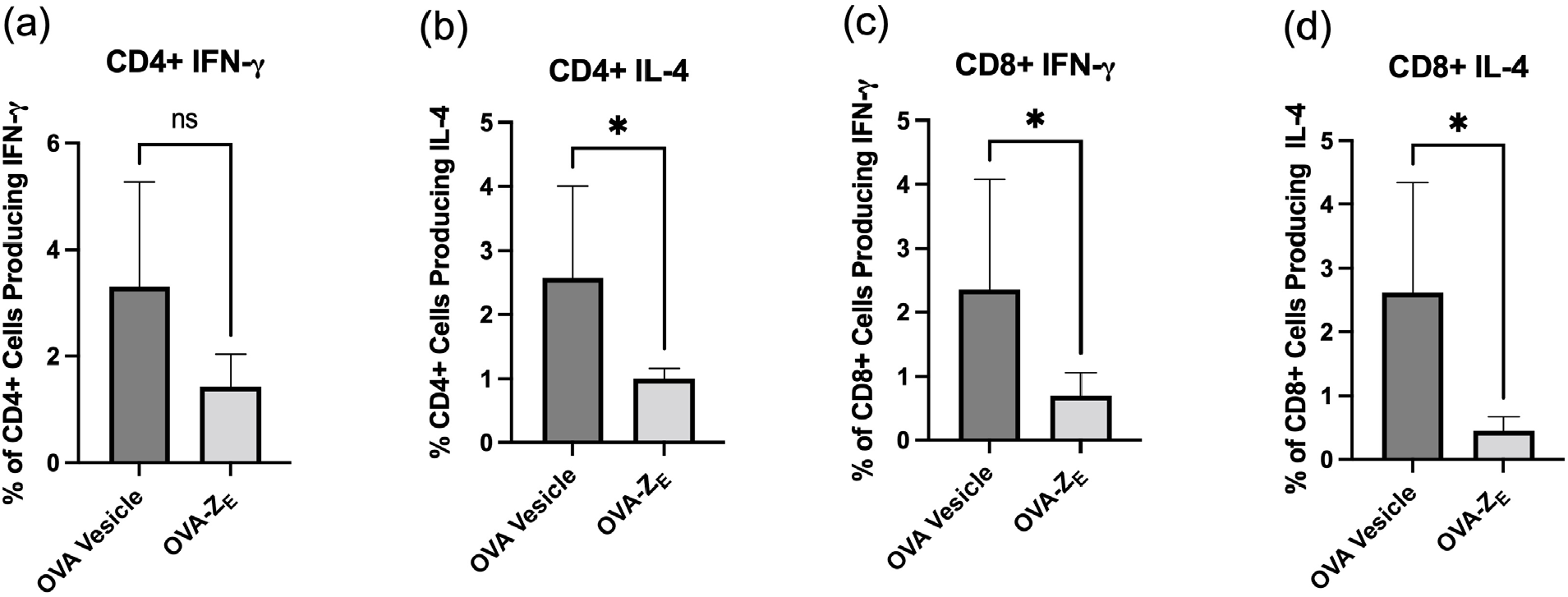
Cytokine production in splenic T cells. Percent of CD4+ cells secreting IFN-γ (a) and IL4 (b). Percent of CD8+ cells secreting IFN-γ (c) and IL4 (d). (*p<0.05)

### 3.6 OVA-Specific T Cell Responses

Higher IgG2a antibody titers induced by the vesicle group suggested there was also a T cell response to OVA protein vesicles [78]. Mice were sacrificed 2 weeks after boost immunization and spleens were collected to analyze OVA-specific T cell responses. Isolated splenocytes were stimulated with OVA and production of IFN-γ and IL4 cytokines by CD4+ and CD8+ splenocytes was measured. The OVA protein vesicle group produced significantly higher amounts of IL4 in CD4+ T cells and both IFN-γ and IL4 in CD8+ T cells than the soluble OVA-Z_E_group. As indicated by the *in vitro* DC maturation study, OVA protein vesicles may have enhanced DC maturation compared to soluble OVA-Z_E_*in vivo* as well. OVA antigens would then be presented via MHC class I and MHC class II molecules in APCs. MHC class II molecules on APCs activated naïve CD4+ T cells and MHC class I molecules stimulated naïve CD8+ T cells. In T cell subsets of splenocytes, Type 2 T helper (Th2) cell is associated with IL4 production, and Th1 is associated with IFN-γ production [79]. Th2 cytokine IL4 promotes B cells to produce IgG1 but inhibits IgG2a production, while Th1 cytokine IFN-*γ* enhances IgG2a production but inhibits IgG1. As shown in our data, significant higher levels of IL4 in CD4+ cells corresponded with higher IgG1 antibody titer. Both humoral and cellular immune responses indicated that OVA protein vesicles induced Th2-biased responses. This is expected for OVA, which is an allergen not a pathogen and elicits a Th2 response when presented in its native conformation [80, 81]. OVA conjugated onto the surface of micelles was reported to induce production of high IgG1 titer with relatively low IgG2a titer [82].

## 4. Conclusion

This work demonstrates the potential of protein vesicles as a subunit vaccine delivery platform. OVA protein vesicles were self-assembled from antigen fusion protein, OVA-Z_E_, and thermoresponsive, photocrosslinkable protein, pZ_R_-ELP. Protein vesicles showed the ability to display full-size antigens on the surface while maintaining the ability to self-assemble, which has not commonly been seen in other self-assembling systems. To our knowledge, this is the first example of using ELP-based self-assembling system to deliver whole antigen proteins. Immunization of mice proved that OVA protein vesicles induced OVA-specific humoral and cellular immune responses appropriate for conformational OVA allergen. Additionally, OVA vesicle stability outside the cold chain could be valuable for translation to all communities. Altogether, the protein vesicle is a promising vaccine platform given its ability to display antigen proteins, long-term stability, tunable size and multivalency, combined with its vaccination efficacy in mice. Previous work has demonstrated that protein vesicles can be made from globular proteins with a wide range of size and surface charge [54]. In future work, antigen proteins from pathogens will be incorporated into protein vesicles to examine the efficiency of protein vesicles to protect again infectious diseases, including simultaneous presentation of multiple antigens. The ability to control antigen density could enable investigation of fundamental questions regarding the effect of multivalent antigen density and spacing on BCR binding and activation. Given the high positive charge in Z_R_and hydrophobic lumen with cargo capacity [42], incorporation of nucleic acid or small molecule adjuvants may be possible to enhance immunogenicity of antigens or bias the nature of the immune response. The modularity and simplicity of protein vesicles could enable a variety of vaccine applications.

## Acknowledgements

The authors acknowledge financial support from the National Science Foundation BMAT Award 2104734. This work was performed in part at the Georgia Tech Institute for Electronics and Nanotechnology, a member of the National Nanotechnology Coordinated Infrastructure, which is supported by the National Science Foundation (Grant No. ECCS-2025462). The authors gratefully acknowledge Prof. D.A. Tirrell and Prof. K. Zhang for Z_E_and Z_R_-ELP genes and AF-IQ *E. coli*. We acknowledge the contributions of named and unnamed people whose health, lives, livelihoods, legacy, and privacy were extorted, often without compensation, consent, or regard to their safety, in the name of biomedical research. These men, women, and children were stripped of their humanity, and often their identity. We knowingly use resources and knowledge with the gratitude and respect not given previously. We commit to educating ourselves and others on the history and ethical failures of biomedical research, expressing our gratitude, and encouraging others to do the same.

## Author contributions

Y.L.: Conceptualization; Formal analysis; Investigation; Methodology; Roles/Writing - original draft; and Writing - review & editing. J.A.C.: Conceptualization; Funding acquisition; Project administration; Supervision; Writing - review & editing.

