## Supplemental Information for "Self-Assembled Protein Vesicles as Vaccine Delivery Platform to Enhance Antigen-Specific Immune Responses"

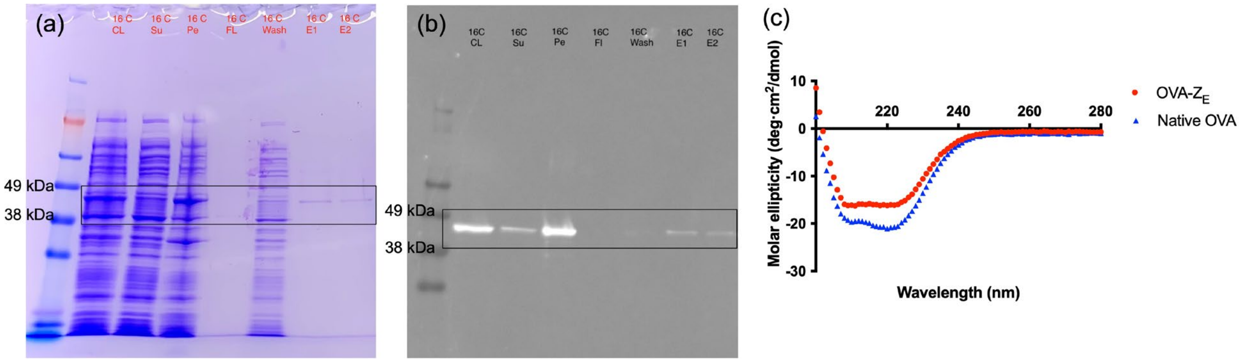


Figure S1. SDS-PAGE gel of OVA-$Z_E$ (a) and western blot image of OVA-$Z_E$ (b). The MW of OVA-ZE is 50.2 kDa. From left to right were products collected from CL: cell lysate (total protein), Su: supernatant of cleared lysate (soluble protein used for purification), Pe: pellet from cleared lysate (insoluble protein), Ft: flow through from Ni-NTA column, Wash, and E1, E2: elutions from Ni-NTA column following protein purification. (c) CD spectra of native (purchased) OVA and recombinant OVA-$Z_E$.

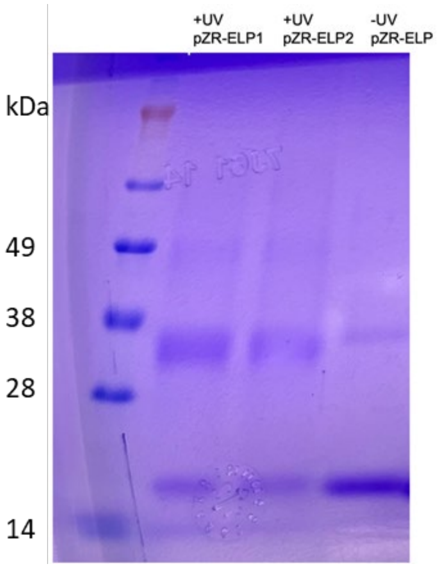


Figure S2. SDS-PAGE gel of $pZ_R$-ELP. Without UV crosslinking (-UV) protein is mainly monomeric (17 kDa) with a small fraction of dimer due to some unreduced disulfide bonds between terminal cysteines. With crosslinking (+UV), there is less monomer, more dimer and some trimer and smeared higher order oligomers due to covalent bonds formed between the proteins. (a) and western blot image of OVA-$Z_E$ (b). The MW of OVA-ZE is 50.2 kDa. From left to right were products collected from CL: cell lysate (total protein), Su: supernatant of cleared lysate (soluble protein used for purification), Pe: pellet from cleared lysate (insoluble protein), Ft: flow through from Ni-NTA column, Wash, and E1, E2: elutions from Ni-NTA column following protein purification. (c) CD spectra of native (purchased) OVA and recombinant OVA-$Z_E$.

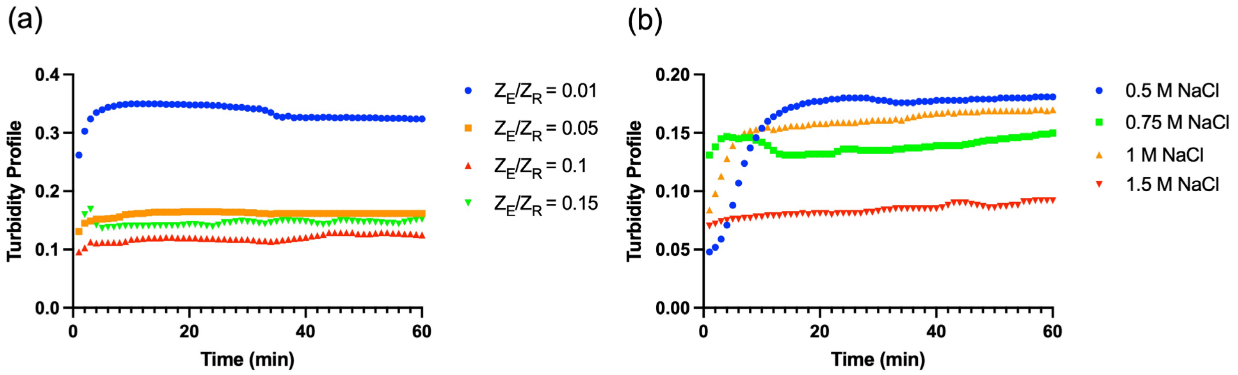


Figure S3. (a) Turbidity profiles (optical density at 400 nm, arbitrary units) of OVA-$Z_E$/$pZ_R$-ELP mixtures at $Z_E/Z_R$ molar ratios of 0.01, 0.05, 0.1 and 0.15 with $pZ_R$-ELP concentration of 30 μM and NaCl concentration of 1 M. (b) Turbidity profile of OVA-$Z_E$/$pZ_R$-ELP mixtures at 0.5 M, 0.75 M, 1 M and 1.5 M NaCl concentrations with 30 μM $pZ_R$-ELP and 3 μM OVA-$Z_E$.

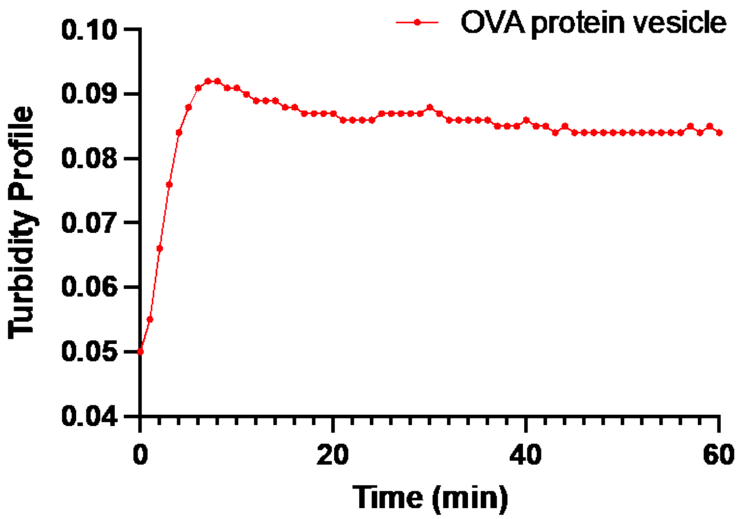


Figure S4. Turbidity profile (optical density at 400 nm, arbitrary units) of OVA-$Z_E$/p$Z_R$-ELP complex containing 3 µM OVA-$Z_E$ and 30 µM p$Z_R$-ELP in 1 M NaCl phosphate buffer. Turbidity profile was measured at 400 nm at 25 °C.

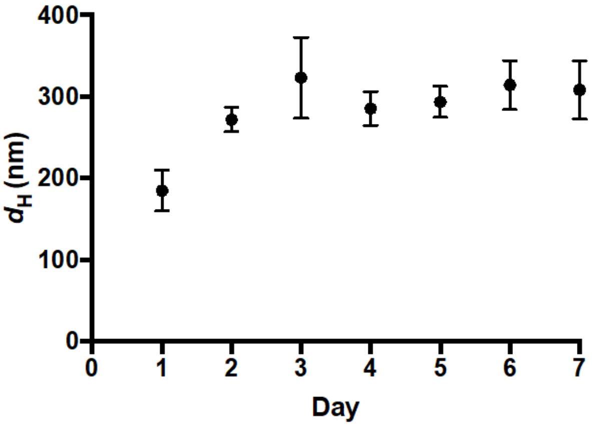


Figure S5. Hydrodynamic diameters of OVA protein vesicles over 7 days after dialysis into PBS as measured by DLS. Each data point is the average of three replicate batches of vesicles.

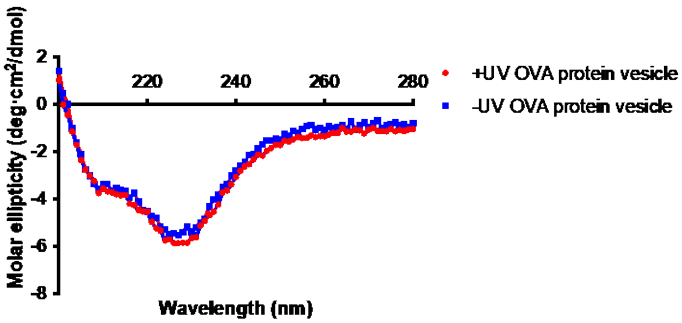


Figure S6. CD spectra of OVA protein vesicles with (red) and without (blue) UV irradiation.
